# Two-locus CRISPR toxin-antidote gene drive for confined population modification

**DOI:** 10.64898/2026.07.12.738060

**Authors:** Ruobing Feng, Andrea Y.N. Tan, Zhuoran Lu, Yuchang Chen, Jackson Champer

## Abstract

Gene drive systems enable rapid spread of desired transgenes throughout populations. Advances in CRISPR technology have facilitated the construction of toxin-antidote gene drives, which utilize a CRISPR nuclease as the toxin to disrupt an essential wild-type gene alongside a recoded version of the same gene as the antidote. Because these systems propagate by eliminating wild-type alleles rather than directly copying themselves like homing drives, they typically exhibit introduction thresholds, allowing them to be confined to target populations. Previous work developed the efficient Toxin-Antidote Recessive Embryo (TARE) drive, but its threshold may be too low in challenging confinement scenarios. Here, we constructed a 2-locus TARE drive system. It has underdominance characteristics, yielding a higher introduction threshold, even when drive performance is ideal. It targets the essential but haplosufficient genes *hairy* and *sim* using two different drives at different genomic locations, each targeting the gene that the other rescues. Our system involved two linked elements together with rare homology-directed repair-mediated drive conversion, reducing the threshold to compensate for fitness costs. The system showed high efficiency in individual crosses. When released into multigenerational cage populations above the introduction threshold, the drive successfully and rapidly modified the entire population, and when below this threshold, it was eliminated. Our findings indicate that 2-locus TARE drives represent promising tools for effective and strongly confined population modification.

## Introduction

Gene drive systems are engineered genetic elements capable of spreading through a population due to their super-Mendelian inheritance (Bier, 2022; Hay et al., 2021; Verkuijl et al., 2022). Even following a modest release of gene drive individuals, the drive can sweep through a population in just a few generations, provided that drive performance is sufficient. This capacity for efficient self-propagation offers potential solutions to global challenges in public health, agriculture, and the environment (Bier, 2022; Hay et al., 2021; Verkuijl et al., 2022; Wang et al., 2024; Yadav et al., 2025). Compared to currently deployed control technologies, such as pesticides, gene drives are highly species-specific (Debrah et al., 2025). Furthermore, they are potentially more cost-effective than other genetic strategies, such as the sterile insect technique (SIT), release of insects carrying a dominant lethal gene (RIDL), and its female-specific (fsRIDL) variant (Fu et al., 2010; Han and Champer, 2025; Thomas et al., 2000) due to the relatively small release sizes required.

Gene drive systems are divided into two categories based on their ultimate effects on the target population (Wang et al., 2022). The first is population suppression drives, which aim to reduce or eliminate harmful populations by targeting genes essential for reproduction, sex determination, or survival, thereby lowering population density (Esvelt and Gemmell, 2017; Hammond et al., 2016). In contrast, population modification drives do not aim to reduce population size but instead rely on the high inheritance efficiency of gene drives to spread specific “cargo genes” or mutations throughout a population, conferring desirable traits such as resistance to pathogens like *Plasmodium* or Dengue virus in mosquito vectors (Hoermann et al., 2022; Li et al., 2025). Collectively, these two strategies provide a powerful toolkit for addressing pressing global issues, including the control of agricultural pests like the diamondback moth (*Plutella xylostella*) (Xu et al., 2022), the management of invasive species, and the mitigation of vector-borne diseases such as malaria, dengue, Zika, chikungunya, and yellow fever (Gubler, 2011; Tahir et al., 2019; Wang et al., 2024; Weaver and Reisen, 2010).

The most intensively researched systems are CRISPR-based homing drives(Verkuijl et al., 2022; Wang et al., 2024), which function by cutting the wild-type chromosome and using the drive-carrying chromosome as a template for homology-directed repair (HDR), effectively converting heterozygotes to homozygotes in the germline (Champer et al., 2017). While powerful, the efficacy of homing drives is often compromised by the formation of resistance alleles. When the DNA break is repaired by the end-joining pathway, small insertions or deletions are created at the target site, preventing further recognition by the drive’s guide RNAs and stopping its spread. Though the worst effects of resistance alleles can be mitigated with guide RNA (gRNA) multiplexing (Anderson et al., 2024; Champer et al., 2018; Champer et al., 2020e; Chen et al., 2025; Khatri and Burt, 2022) (plus rescue elements for modification drives), they can still contribute to reduced efficiency. Furthermore, homing drives are typically invasive systems, where the release of even a few individuals can lead to the drive spreading to any connected population, raising ecological and sociopolitical concerns where spatial confinement is desired (Rode et al., 2020).

Toxin-Antidote gene drives offer a promising solution to the dual challenges of resistance and confinement. They function by eliminating wild-type alleles. A typical CRISPR-based Toxin-Antidote system consists of a “toxin” (Cas9 and gRNAs targeting an essential gene) and an “antidote” (a recoded, functional version of the target gene immune to cleavage) (Champer et al., 2020c; Oberhofer et al., 2019). Unlike homing drives, which convert wild-type alleles into drive alleles only through HDR, toxin-antidote systems convert wild-type alleles into nonfunctional forms through either end-joining or HDR. By leveraging the formation of these disrupted alleles, the toxin-antidote mechanism offers broad potential for application across diverse organisms.

HDR in toxin-antidote systems can produce several distinct outcomes depending on the extent of homology used during repair. In a favorable outcome, repair copies only the local sequence disruptions adjacent to the drive allele, producing a nonfunctional target allele. This has a similar effect to end-joining but is less likely to preserve a functional target sequence. A rarer long-range HDR event may copy the entire drive allele, creating a drive conversion event that can increase drive inheritance (Champer et al., 2020c). Conversely, partial HDR that copies only the recoded rescue sequence without the full drive cassette could generate a functional resistance allele, which would negatively affect drive success. Previous TARE designs therefore considered the relative likelihood of local disruptive repair, full-drive copying, and recoded-sequence-only repair when placing gRNA target and rescue sequences, allowing for substantial DNA between the drive insertion site and target, and between the target site and 3′ UTR of the target gene (Champer et al., 2020c).

CRISPR toxin-antidote gene drives have many variants based on the attributes of the target genes and the arrangement of drive components (Champer et al., 2020a; Champer et al., 2020b). When the target gene is haplosufficient, a Toxin-Antidote Recessive Embryo (TARE) drive is formed, causing nonviability in individuals that inherit two disrupted alleles (Champer et al., 2020c; Metzloff et al., 2022; Oberhofer et al., 2019, 2020). When the essential gene is haplolethal, a Toxin-Antidote Dominant Embryo (TADE) drive is formed, in which even a single disrupted wild-type allele is lethal (Champer et al., 2020a; Champer et al., 2020b).

In terms of the arrangement of drive components, systems can be classified as either single-locus or multi-locus (Champer et al., 2020a). A single-locus system can utilize either a single drive allele that targets and provides rescue for the same gene, or two distinct drive alleles where each targets the gene rescued by the other. Examples of such designs include the same-site TARE or the distant-site Cleave and Rescue (ClvR) drive. The engineering trade-off between same-site designs and distant-site configurations involves the balance between expression fidelity and targeting flexibility. Same-site drives facilitate rescue expression by utilizing endogenous regulatory sequences (Chen et al., 2023), whereas distant-site drives offer expanded target site selection and higher tolerance for low cutting rates. Multi-locus systems place different drive alleles at separate genetic sites. Because these different combinations of target gene and component arrangement result in different removal ratios of wild-type versus drive alleles, they allow for the fine-tuning of a drive’s population dynamics, producing systems with a wide range of introduction thresholds and rates of spread. In multi-locus drive systems, the distance between component loci can also shape drive dynamics. If two drive components are tightly genetically linked, they may be co-inherited, increasing the probability that the complete drive system is transmitted together. Conversely, recombination between the loci can separate the components, generating single-drive haplotypes and reducing the efficiency of establishment. Thus, linkage and recombination provide an additional layer of control over the introduction threshold, especially for drive systems whose spread depends on the joint inheritance of multiple drive elements.

The “introduction threshold” is the key to creating confined gene drives. It establishes a minimum frequency of drive-carrying individuals required for the drive to spread; if introduced below this level, the drive is naturally eliminated. This property provides two major benefits: spatial confinement, which can restrict a drive to a target area, and reversibility, as a release of wild-type individuals can dilute the drive’s frequency below the threshold (Champer et al., 2020d; Edgington et al., 2020; Kläy et al., 2025). However, the degree of confinement is not absolute and depends on ecological factors such as migration rates between populations. For a drive to remain localized, the rate of migration to adjacent non-target populations must remain below a critical “migration threshold”. The migration threshold is correlated with the introduction threshold. It represents the maximum migration rate that can be sustained before steady influx of drive alleles in a new population overcomes drive loss, pushing the drive above its introduction threshold and resulting in full spread of the drive into the new population. Considering only individuals with two disrupted wild-type alleles will be removed from the population, a TARE drive will not have introduction threshold unless it has a fitness cost (Champer et al., 2020a; Champer et al., 2020b). While at least small fitness costs are likely in real systems, the low resulting threshold may be insufficient for some applications requiring stringent confinement. Recent ecological modeling suggests that the invasion threshold for underdominance drives is an intrinsic property governed by drive performance, largely independent of ecological factors like density-dependent feedback (Xu and Bonsall, 2025). Consequently, environmental context cannot mitigate such design imperfections or artificially elevate a low threshold. Thus, it is important to carefully select a drive with a specific threshold for a specific application.

Some toxin-antidote systems can be configured for underdominance, a genetic phenomenon proposed early as a gene drive mechanism (Curtis, 1968; Magori and Gould, 2006; Reeves et al., 2014; Serebrovsky, 1940) where heterozygotes possess lower fitness than either type of homozygote. One possible architecture is the 2-locus TARE drive. In this system, two TARE elements are placed at separate genetic loci, each rescuing the gene at their locus and targeting the wild-type allele at the other site for disruption. This creates a mutual-dependency system where only individuals with both drive elements can consistently survive. Accordingly, some of the drive elements will be removed from the population alongside with the removal of disrupted alleles, resulting in an introduction threshold of approximately 18% in the absence of fitness costs (Champer et al., 2020a). This built-in confinement makes the 2-locus TARE system a highly attractive candidate over a range of drive performance parameters.

Because the 2-locus TARE system depends on the co-inheritance of two drive components, its dynamics may be strongly affected by genetic linkage. Tight linkage is expected to facilitate establishment, whereas recombination can break this association and generate single-drive haplotypes, thereby increasing the introduction threshold. Further, HDR-mediated drive conversion may restore or amplify drive alleles and partially counteract recombination-mediated drive allele separation. These features suggest that linkage, recombination, and HDR could be used to tune the confinement properties of a 2-locus TARE drive.

While the theoretical advantages of 2-locus TARE drive are compelling, its performance has yet to be empirically demonstrated. In this study, we report the construction and experimental validation of an asymmetric HDR-assisted linked 2-locus TARE gene drive in *Drosophila melanogaster*. We selected two haplosufficient essential genes, *hairy* and *single-minded* (*sim*), located on the same chromosome, allowing us to examine how genetic linkage and sex-specific recombination influence drive establishment. In addition, we incorporated an asymmetric gRNA design that allows HDR-mediated conversion to act as a threshold-modulating process rather than as the primary drive mechanism. We assessed its drive inheritance and performance with individual crosses, and also evaluate its capacity to spread through large cage populations following releases of different sizes. Our results provide an experimental demonstration of a confined, 2-locus TARE system and show how genetic linkage and HDR can jointly tune its introduction threshold.

## Methods

### Plasmid construction

To generate the final constructs for injection, 2LTAREsimNUh4G (expressing a DsRed marker) and 2LTAREhNUsim4G (expressing an EGFP marker) were created via Gibson assembly. All reagents used for PCR, Gibson assembly, and plasmid miniprep were purchased from New England Biolabs and Vazyme. Oligonucleotide primers were synthesized by Tsingke Biotechnology. 5-α competent Escherichia coli cells were obtained from Vazyme, and plasmid midipreps were performed using the ZymoPure Midiprep Kit (Zymo Research). All constructed plasmids were verified by Sanger sequencing. All drive and injection plasmids are available on GitHub (https://github.com/jchamper/Two-locus-TARE-Drive).

### Generation of transgenic lines

Embryo injections were performed by Unihuaii Company. Donor plasmids 2LTAREsimNUh4G and 2LTAREhNUsim4G (500 ng/μL) were co-injected into *w^1118^*embryos along with TTChsp70c9 (500 ng/μL) and the gRNA plasmids (100 ng/μL), TTTh4g, TTTsim2g, providing the Cas9 protein and guide RNAs required for transformation. Two transgenic lines were generated. In one line, the drive element was inserted into the *hairy* gene and targeted the *sim* gene, and these flies exhibited DsRed fluorescence in the eyes. In the other line, the drive element was inserted into the *sim* gene and targeted the *hairy* gene, and these flies exhibited EGFP fluorescence in the eyes. Injected males displaying either DsRed or EGFP fluorescence, which indicated successful drive insertion, were crossed with *w^1118^*flies to establish these single-locus drive lines. Next, individuals with different fluorescent markers were crossed to generate heterozygous lines expressing double fluorescence. These flies were then intercrossed over several generations to establish homozygous lines. Preference was given to individuals displaying noticeably brighter eye fluorescence, which typically suggests homozygosity. Finally, the homozygous status of flies carrying both DsRed and EGFP markers was confirmed by individual crosses with *w^1118^* flies.

### Fly Rearing and Phenotyping

Flies were maintained in an incubator at 25°C under a 14:10 h light/dark cycle with 60% relative humidity. For phenotyping, adult flies were anesthetized using CO₂ and screened for DsRed and EGFP fluorescence using the NIGHTSEA illumination system.

### Drive efficiency and egg viability test

We assessed the drive efficiency of individuals with drives at one locus and individuals with drive alleles at both loci. For the single-locus drive performance test, drive heterozygous lines were generated from crosses between fluorescent males and *w^1118^* females. For the double-loci drive performance and inheritance test, drive homozygous lines were crossed to the *w^1118^* line to generate drive-heterozygous offspring. These heterozygous drive offspring were then crossed either to *w^1118^*flies or to their own siblings. All crosses were consistently set up using two males and two virgin females in each vial. The resulting progeny were screened for fluorescence expression to determine their genotype and quantify drive efficiency.

At the same time, we assessed the egg viability of heterozygous drive individuals with either one drive allele type or one of each type. Eggs laid from these crosses were collected and counted daily on three successive days at 24-hour intervals. After development, the number of emerged adults was counted. To assess nonviability associated with the drive elements, the egg-to-adult viability of drive-carrying individuals was compared to that of the *w^1118^* control and also with each other.

We processed our drive efficiency and egg viability data using a generalized linear mixed-effects model (GLMM) following a binomial distribution. Because each experimental vial acts as a distinct batch, ignoring this structure can artificially inflate statistical confidence. To address this, we incorporated the vial identifier as a random effect within the model, utilizing Adaptive Gauss-Hermite Quadrature (maximum likelihood, nAGQ = 25) to accurately capture variance and yield robust standard errors. The analyses were conducted in R (version 4.1.0) to extract the estimated marginal means from these models. Finally, we employed z-tests on these estimates to execute pairwise group comparisons and to compare to the baseline viability of the *w^1118^* strain.

### Recombination assay

We estimated the recombination rate between the *hairy* and *sim* loci using two non-driving DsRed marker lines in *hairy* and *sim*. Double-heterozygous flies were generated by crossing *hairy* line homozygotes to *sim* line homozygotes, which were then crossed to *w^1118^* flies. Each vial contained one male and one female. The total recombination rate was therefore calculated as twice the proportion of non-fluorescent progeny.

Recombination frequency was analyzed using a generalized linear mixed-effects model (GLMM) with a binomial error distribution. Because each experimental vial represented a distinct replicate, vial identity was included as a random effect to account for vial-to-vial variation and avoid inflated statistical confidence. Models were fitted by maximum likelihood using adaptive Gauss–Hermite quadrature (maximum likelihood, nAGQ = 25). Analyses were conducted in in R (version 4.1.0), and estimated marginal means of the non-fluorescent progeny proportion were extracted on the response scale. The final sex-specific recombination rate was obtained by multiplying this estimated proportion by two.

### Quantitative modeling of cut rates

To estimate the molecular efficiency of our TARE drive system, we developed a quantitative model based on observed egg-to-adult viability and drive inheritance data. We assumed that the reduced viability observed in drive crosses, relative to wild-type control crosses, was caused by the formation of disrupted recessive lethal alleles through Cas9-mediated cleavage at the target loci. For each locus, lethality was governed by two parameters: the effective germline disruption probability, 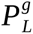, defined as the probability that a gamete carries a disrupted allele after germline cleavage, and the embryo cleavage rate, 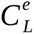, defined as the probability that a wild-type allele is cleaved by maternally deposited Cas9/gRNA in the early zygote.

We jointly estimated four parameters, 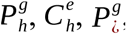, and 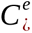, where the subscripts denote the *hairy*-associated and *sim*-associated target loci, and the superscripts *g* and *e* denote germline disruption and embryo cleavage, respectively. Parameter estimation was performed by simultaneously fitting eight equations: four describing egg-to-adult lethality and four describing drive inheritance. This joint fitting strategy allowed the cleavage parameters to be constrained by both progeny survival and the observed transmission of each drive allele.

For a single-locus drive cross between a heterozygous female and a wild-type male, the frequency of lethal progeny at locus *L*, where *L* ∈ {*h*, ∼}, was modeled as:

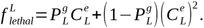

The first term represents the probability that a progeny inherits a disrupted allele from the mother and that its paternal wild-type allele is cleaved by maternally deposited Cas9/gRNA. The second term represents the probability that the progeny inherits a maternal wild-type allele, which is subsequently cleaved in the embryo together with the paternal wild-type allele. This single-locus equation was applied separately to the *hairy*-associated and *sim*-associated single-drive crosses, yielding the first two equations in the joint model.

For double-locus crosses, we explicitly modeled the phased two-locus gamete distribution rather than treating the two loci as independent marginal genotypes. Throughout this section, subscripts *h* and ¿ denote the allelic states or target loci, whereas superscripts *g* and *e* denote germline and embryo processes. Let *q_f_* (*g_h_, g*_¿_) denote the probability that a female gamete carries allele *g_h_* at the hairy-associated locus and allele *g*_¿_ at the *sim*-associated locus after germline cleavage and recombination. Similarly, let *q_m_* (*p_h_, p*_¿_) denote the corresponding male gamete distribution. Here, *g_h_, p_h_* ∈ {*D_h_, R_h_, W_h_* } and *g*_¿_, *p*_¿_ ∈ {*D*_¿_, *R*_¿_, *W* _¿_ }, where *D*, *R*, and *W* represent drive, disrupted resistant, and wild-type alleles, respectively. Thus, each gamete belongs to one of nine possible phased two-locus haplotypes. This phased representation allowed recombination to be incorporated using the experimentally specified sex-specific recombination rates, with *r_f_* for females and *r_m_* for males. In this formulation, *q_f_* and *q_m_* are not independently fitted parameters, but deterministic gamete distributions calculated from the germline disruption probabilities 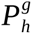 and 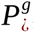, together with the fixed sex-specific recombination rates. The embryo cleavage parameters 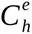 and 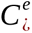 enter the model through the per-locus survival functions.

For a double-heterozygous female crossed to a wild-type male, the survival probability was calculated as:

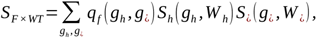

and the corresponding lethal fraction was:

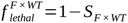

For each locus *L*, the per-locus survival function in this cross was defined as:

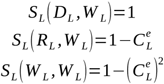

Thus, progeny carrying a drive allele were protected by the rescue sequence. Progeny inheriting a disrupted allele required cleavage of the paternal wild-type allele to become lethal, whereas progeny inheriting a maternal wild-type allele required cleavage of both maternal and paternal wild-type alleles. Overall lethality was derived by subtracting the joint two-locus survival probability from one.

For a double-locus drive cross between double-heterozygous females and double-heterozygous males, the survival probability was calculated as:

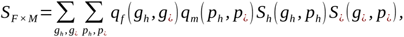

and the corresponding lethal fraction was:

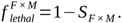

For each locus *L*, the per-locus survival function was defined as:

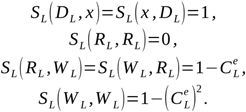

Unlike the female-to-wild-type cross, both parents contribute gametes that may carry disrupted alleles generated through germline cleavage. Therefore, lethality in the double-heterozygous female-by-male cross was calculated by summing over all maternal and paternal phased gamete combinations, rather than by multiplying independent single-locus marginal probabilities.

Drive inheritance was modeled conditional on progeny survival. Here, 1() denotes an indicator function, which equals 1 if the progeny carries the specified drive allele and 0 otherwise. For the double-heterozygous female crossed to wild-type males, inheritance of the hairy-associated drive was modeled as:

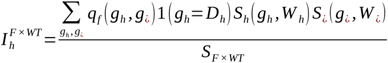

and inheritance of the sim-associated drive was modeled as:

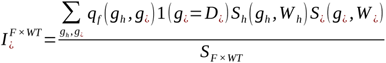

For the double-heterozygous female-by-male cross, drive inheritance was calculated as the probability that a surviving progeny carried at least one copy of the corresponding drive allele. Thus, inheritance of the *hairy*-associated drive was modeled as:

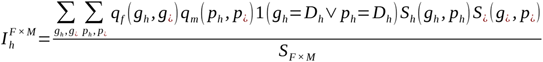

and inheritance of the *sim*-associated drive was modeled as:

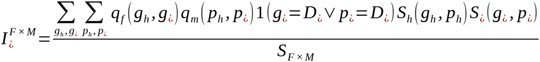

Together, the model fitted eight observed quantities: single-locus lethality for the *hairy*-associated drive, single-locus lethality for the *sim*-associated drive, double-locus lethality in the female-by-wild-type cross, double-locus lethality in the female-by-male cross, *hairy*-associated drive inheritance in the female-by-wild-type cross, *sim*-associated drive inheritance in the female-by-wild-type cross, *hairy*-associated drive inheritance in the female-by-male cross, and sim-associated drive inheritance in the female-by-male cross.

We used a weighted non-linear least-squares approach to fit the experimental viability and drive inheritance data to the eight equations described above. The best-fit parameter set was selected by minimizing the weighted percentage error across all eight equations. A grid search was performed across the four parameters 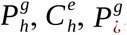, and 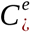, with the constraint that the effective germline disruption probability exceeded the embryo cleavage rate at each locus.

In addition, we imposed a biological boundary constraint to exclude parameter combinations in which both germline disruption probabilities simultaneously reached complete disruption, that is, 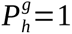 and 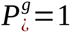. This constraint was motivated by the drive inheritance data from double-locus crosses. Although drive inheritance was high, the observed inheritance of the two drives in double-heterozygous female-by-male crosses did not reach 100% (Figure 4A), indicating that both loci were unlikely to have complete effective germline disruption simultaneously under the experimental conditions. Therefore, parameter combinations with 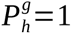 and 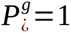 were excluded from the search space.

### Cage study

Flies were maintained in 30 × 30 × 30 cm enclosures containing nine food bottles, with the oldest bottle replaced approximately every 12-14 days. Double homozygous drive flies and age-matched *w^1118^* flies were then mixed at drive-to-wild-type ratios of 12%, 24%, 36%, 40%, and 48%, distributed evenly into nine separate food bottles, and allowed to lay eggs for one day before being removed. The bottles were subsequently placed into cages, and the emerging flies were designated as the initial generation. All were either drive homozygotes (at both insertion sites) or wild-type. After 12-14 days, the bottles were replaced with fresh food while the adult flies remained in the cage. One day later, flies were removed from bottles for phenotyping (flies were frozen at this point), and then bottles were returned to the cage. In each subsequent generation, flies were allowed to develop in the cage for 12-14 days (extended past 12 days only if more than ∼50% of pupae had not yet eclosed). New bottles were then introduced into the cages for 24-48 hours to collect eggs for the next generation (extended to 48 hours only if the number of eggs was substantially lower than in the previous generation).

### Estimation of drive fitness

We used a maximum likelihood approach to estimate fitness parameters for the gene drive lines. Details of this approach and its applications to cage population data were previously described(Liu et al., 2019). For the two-locus TARE line, we estimated the viability, female fecundity, and male mating success parameters and the effective population size using a model that assumes a multiplicative fitness effect of each TARE allele (homozygotes were assigned fitness equal to the square of heterozygotes). Both loci are haplosufficient, so heterozygotes that bear one wild-type and one disrupted allele were assumed to have the same fitness as wildtype homozygotes. Fitness effects were assumed to be multiplicative across loci. Resistance homozygotes at either site were modeled as recessive lethal.

The germline and embryo rates of cleavage of the wild-type allele by TARE were set at experimentally estimated values. The rare drive conversion rate and the sex-specific recombination rate between *hairy* and *sim* were also derived from experimental data. Parameter values were estimated by maximizing the likelihood across all generation transitions from the five cages combined (i.e., a single estimate per parameter is generated for all generations of all five cages combined together). The effective population size parameter was estimated as a fraction of the census population size in each generation transition, using the average of both generations that were part of the transition.

### Simulations

Predictive simulations were conducted using the forward-in-time, individual-based framework SLiM (version 5.0) (Haller et al., 2026), adapting models from previous studies that included 2-locus TARE (Pan and Champer, 2023). We extended this framework to model an HDR-and recombination-modulated 2-locus TARE drive. In addition to standard TARE-mediated conversion of wild-type alleles into disrupted alleles, the model included HDR-mediated drive conversion, allowing a cleaved wild-type allele to be converted into a drive allele at a specified rate. Variable recombination between the two drive loci was also explicitly modeled.

These simulations modeled a panmictic population with the following lifecycle rules: First, each fertile female selects a mate with a probability proportional to the male’s fitness. Second, female fecundity is calculated using a Beverton–Holt curve: 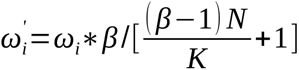, where N is population size, K is carrying capacity, and β is the low-density growth rate. Third, the number of offspring is drawn from a binomial distribution (max 50) that produces an average of 2 when female fecundity fitness is 1 (wild-type) and the population density is at its carrying capacity. Finally, offspring genotypes are determined according to the drive mechanism.

Separately, to analyze the experimental cage data, we implemented a statistical model in R (version 4.1.0). Using the same maximum likelihood estimation logic as our fitness methods, this R framework integrated experimentally derived cleavage rates, HDR rate and recombination rate. This allowed us to generate predicted population trajectories from the inferred fitness parameters and compare them with the observed data. The source code for all simulations and analyses is publicly available on GitHub (https://github.com/jchamper/Two-locus-TARE-Drive).

### Introduction threshold detection and smoothing

Introduction thresholds were estimated from replicate stochastic simulations across a series of initial drive allele frequencies. For each parameter combination, a given initial drive allele frequency was considered above the threshold if the drive allele frequency increased during any ten consecutive generations within the first 50 generations. This establishment criterion was designed to distinguish sustained drive increase from short-term stochastic fluctuations. For each initial frequency, 50 replicate simulations were performed, and the introduction threshold was defined as the lowest release frequency at which at least 50% of replicates satisfied this criterion. Threshold searches were conducted over a predefined range of introduction frequencies using the specified introduction-frequency step size. To avoid biologically unrealistic or highly stochastic releases consisting of extremely few drive individuals, the minimum tested introduction frequency was set to 1%. Threshold simulations were performed with a population size of 10,000. For parameter combinations in which the estimated threshold approached the lower search bound of 1%, simulations were repeated with a larger population size of 50,000 to reduce finite-population stochasticity and better resolve near-boundary establishment behavior.

To visualize the effects of recombination and rare HDR on the introduction threshold, one-dimensional threshold scans were smoothed using LOESS regression. For each scan, either recombination rate or rare HDR rate was varied while the other parameter was held constant. Duplicate x-axis values were averaged before smoothing. LOESS smoothing was performed using a second-degree local polynomial with a span of 0.50, and smoothed threshold values were predicted over 300 evenly spaced grid points. To avoid extrapolation artifacts, smoothed values were constrained within the observed range of simulated thresholds. In parallel, a monotone-smoothed curve was generated using isotonic regression followed by monotone spline interpolation, providing a trend line that preserves the expected directional relationship between the varied parameter and the introduction threshold.

## Results

### 2-locus TARE drive mechanism and population dynamics

The two-locus toxin-antidote recessive embryo (TARE) drive is a reciprocal-rescue gene drive architecture designed to bias inheritance while maintaining threshold-dependent spread. In the originally described form of this system, two distinct drive alleles are placed at two different loci, and each drive allele both disrupts an essential gene at the other locus and rescues disruption at its own locus via a recoded, nuclease-resistant rescue sequence. Because the targeted genes are assumed to be essential but haplosufficient, individuals carrying at least one functional (wild-type or drive) copy at each locus remain viable, whereas individuals that lose function at either locus become nonviable.

The fundamental logic of the system is reciprocal targeting and self-rescue (Figure 1A). The drive allele inserted at the first locus encodes a Cas9/gRNA complex (“toxin”) that cleaves the wild-type allele at the second locus, while simultaneously providing a recoded, cleavage-resistant sequence (“antidote”) for rescue its own locus. Conversely, the drive allele at the second locus does the same. This mutual targeting creates a strict co-dependency: the disruption of the essential wild-type gene at one locus can only be compensated for by the specific rescue element provided by the drive allele at the other locus. Consequently, when disrupted alleles become common viability is largely restricted to individuals that possess at least one copy of the drive allele at both loci.

**Figure 1.**
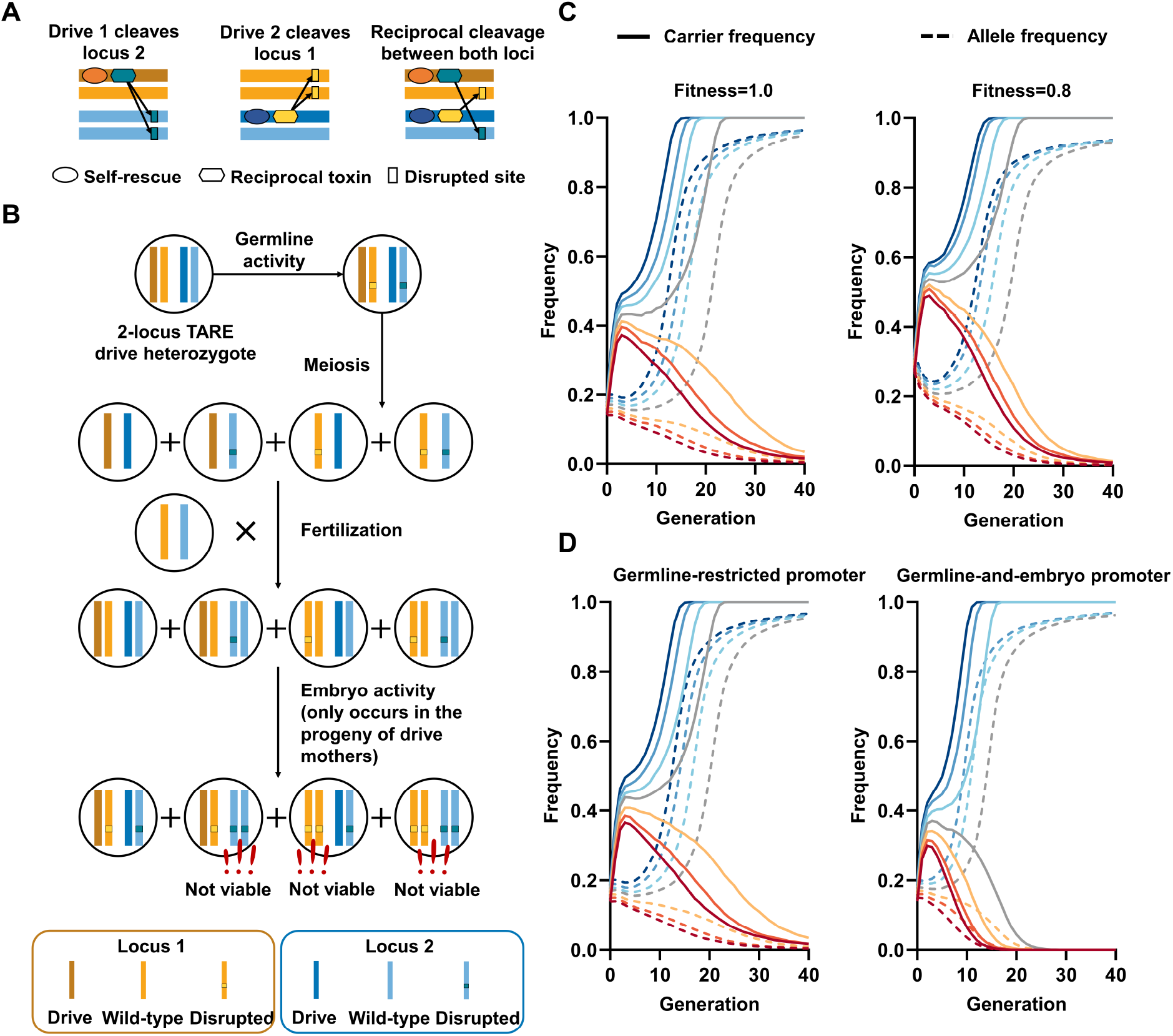
Mechanism and theoretical performance of the canonical two-locus TARE drive. (**A**) The system utilizes two haplosufficient but essential loci. Each drive allele cleaves the wild-type allele at the other locus while providing a rescue element for its own locus. (**B**) The diagram illustrates a cross between a female heterozygous for both drive alleles and a wild-type male. Germline-expressed Cas9/gRNA disrupts wild-type alleles in the oocyte. Maternally deposited Cas9/gRNA complexes disrupt any wild-type alleles in the zygote/early embryo, even if paternally inherited. Viability is contingent upon possessing at least one functional copy (wild-type or drive) at each locus. **(C)** Trajectories of panmictic modeling of 100,000 individuals reveal a bistable outcome. The drive is rapidly eliminated by natural selection when introduced below a specific invasion threshold but sweeps to fixation when released above this frequency. This threshold increases when the drive confers a fitness cost. For the drive with no fitness cost (fitness = 1.0), initial frequencies ranged from 0.14 to 0.20. For the drive with a 20% fitness cost (fitness = 0.8), we tested a higher range of initial frequencies from 0.27 to 0.33. (**D**) As part C, but with a germline-restricted promoter (no cleavage in the early embryo) compared to a promoter that yields 100% cleavage in both the germline and early embryo. For both scenarios, initial release frequencies ranged from 0.14 to 0.20.

This interdependence translates into a distinct inheritance bias at the population level, as illustrated by the drive mechanism (Figure 1B). In a female heterozygous for both drive alleles, Cas9/gRNA complexes target the wild-type *hairy* and *sim* genes in the germline. When she mates with a wild-type male, maternally deposited Cas9 and gRNA persist in the zygote/early embryo, disrupting paternal wild-type alleles. Consequently, assuming 100% cleavage efficiency in both the germline and embryo, survival is restricted to the 25% offspring with both drive alleles. This eliminates wild-type alleles relative to drive alleles at a 5:1 ratio, thereby driving the enrichment of the engineered system among the survivors. Males also contribute to drive spread by disrupting wild-type alleles in the germline, which can later form nonviable genotypes.

This lethality generates distinct population dynamics, characterized by a threshold-dependent spread mechanism (Figure 1C, Data Sets S1-S2). Because the results in removal of both drive and wild-type alleles, it cannot invade from very low initial frequencies. Modeling indicates an invasion threshold of approximately 18% (Champer et al., 2020a). Releases below this critical frequency lead to rapid elimination, whereas releases exceeding the threshold result in the drive’s fixation. Notably, this threshold frequency increases with reduced drive reduced fitness, and fitness costs will also prevent drive fixation (though the drive carrier frequency is still expected to reach 100%).

Building on these fundamental dynamics, we next investigated how specific design choices could further modulate the drive’s behavior. The choice of promoter driving the Cas9 nuclease is important due to the trade-off between the ease of initial invasion and the subsequent speed of population replacement. Promoter performance is primarily governed by the presence or absence of maternally deposited Cas9, leading to cleavage of wild-type alleles in offspring of drive females. To evaluate this, we modeled a germline-restricted promoter (0% embryo cleavage efficiency) and a germline-and-embryo promoter (100% embryo cleavage efficiency). Our simulations revealed that the germline-restricted promoter requires a slightly lower introduction threshold to successfully invade (Figure 1D, Data Sets S1-S2). By confining Cas9 activity strictly to the germline, it avoids removal of drive alleles in some crosses. In contrast, the germline-and-embryo promoter overall cleaves more wild-type alleles and thus facilitates more rapid drive fixation if its slightly higher introduction threshold is exceeded (Figure 1D).

These models establish the basic threshold-dependent behavior of a two-locus TARE drive. However, because successful spread of this system depends on the joint inheritance of both drive alleles, the genetic relationship between the two loci can itself be used as a design parameter. We therefore extended the model architecture into a linked asymmetric variant, in which the two drive components are placed at *hairy* and *sim*, two loci located on the same chromosome. This design allowed us to examine how physical linkage, sex-specific recombination, rare HDR-mediated drive copying, and asymmetric gRNA placement jointly modulate the introduction threshold.

### Design of an asymmetric linked two-locus TARE drive

To build this linked two-locus TARE variant, we engineered drive alleles targeting two distinct genetic loci on chromosome 3 of *Drosophila melanogaster* (Figure 2A-B). The *hairy* gene, which encodes a bHLH transcription factor essential for embryonic segmentation (Carroll et al., 1988), and the *single-minded* (*sim*) gene, a master regulator of central nervous system midline differentiation (FlyBase, FBgn0004666, accessed July 2026) (Nambu et al., 1991), were selected as targets due to their essential yet haplosufficient nature (Champer et al., 2020c; Öztürk-Çolak et al., 2024). In addition to satisfying the biological requirements of TARE targets, these two loci are located on the same chromosome, allowing the *h*-drive and *sim*-drive to be inherited as a partially coupled double-drive haplotype.

**Figure 2.**
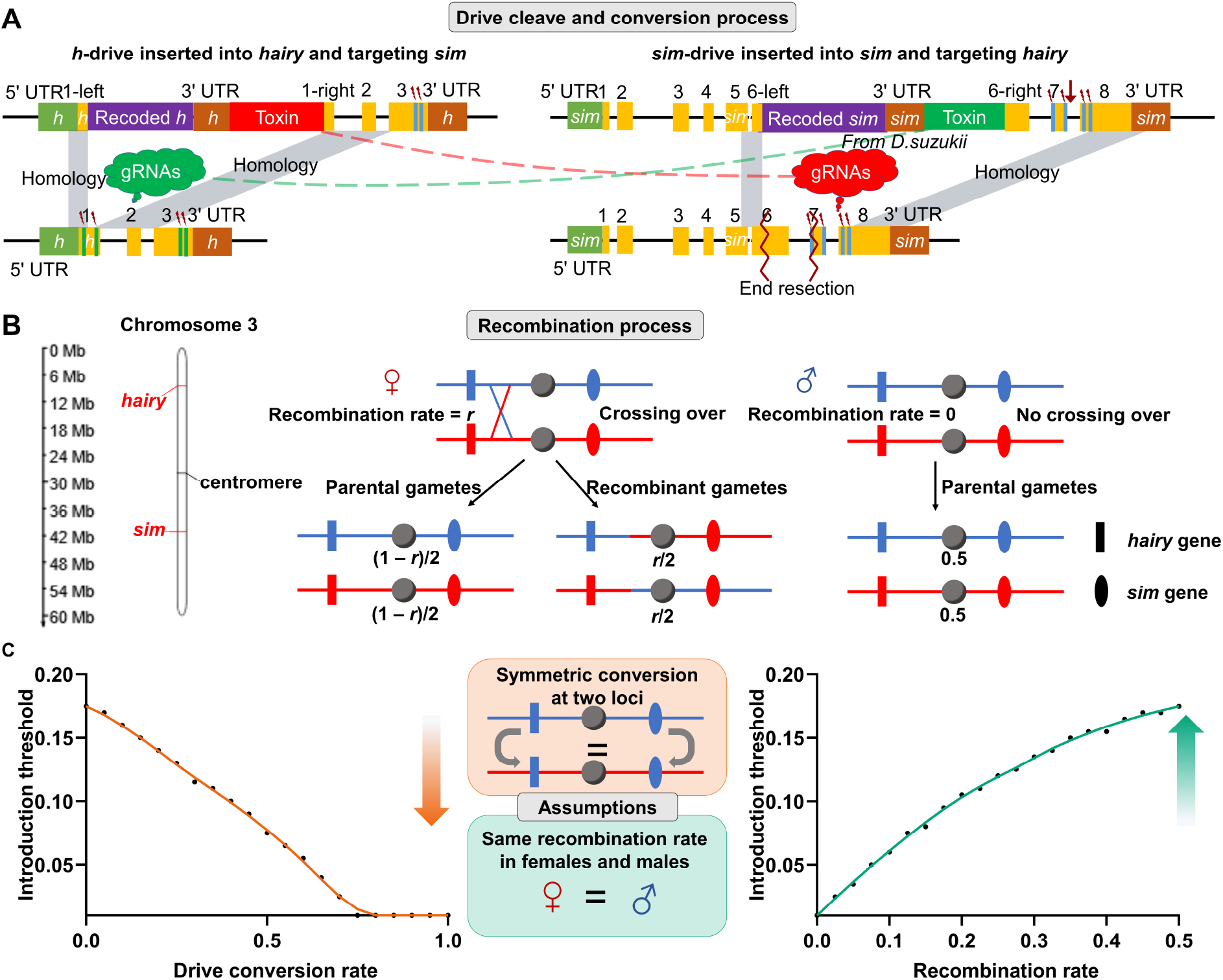
Design of an asymmetric HDR-assisted linked two-locus TARE drive. **(A)** Drive cut and conversion process. The *h*-drive is inserted into *hairy* and targets distal regions of *sim*, functioning as a conventional TARE component that disrupts wild-type *sim* alleles while rescuing *hairy* function through a recoded *hairy* sequence. The *sim*-drive is inserted into *sim* and targets *hairy* through an asymmetric dual-gRNA design. Distal *hairy*-targeting gRNAs function as conventional TARE gRNAs that generate loss-of-function hairy alleles, whereas insertion-proximal gRNAs target the *h*-drive insertion region and provide a potential route for rare HDR-mediated copying of the *h*-drive. **(B)** Recombination process between *hairy* and *sim* on chromosome 3. Because the two loci are located on the same chromosome, the *h*-drive and *sim*-drive are more often inherited together if they start on the same chromosome. In females, recombination occurs between the two loci at rate *r*, generating both parental and recombinant gametes. In males, recombination is absent, preserving the parental linked haplotypes. For the *hairy*-*sim* interval, male recombination was modeled as 0 and female recombination as 30.6%. **(C)** Simulated effects of drive conversion and recombination on the introduction threshold. IN this model, germline and embryo cutting were set to 100%, and no drive alleles carried fitness costs. Drive conversion, representing rare and potentially intended HDR-mediated drive copying, reduces the introduction threshold, whereas recombination increases the threshold by breaking the association between the two drive components. The schematic in the center summarizes the simplifying assumptions used in the parameter scan, including symmetric conversion at the two loci and equal recombination rates between sexes for the scan shown.

This linked arrangement was designed to promote co-inheritance of the complete reciprocal-rescue module while retaining threshold-dependent spread. In *Drosophila melanogaster*, meiotic recombination is absent in males but occurs in females (Vazquez et al., 2002). Therefore, male transmission is expected to preserve the linked *h*-drive/*sim*-drive alleles, whereas female recombination can partially separate the two components and generate single-drive alleles (Figure 2B). Based on our recombination assay, we estimated the recombination rate between *hairy* and *sim* to be 0 in males and 30.6% in females, and we used these values in the model. Because recombination reduces the co-transmission of the two drive components, increasing recombination is expected to raise the introduction threshold.

Both constructs use a polycistronic gRNA design, where individual gRNAs are flanked by tRNA sequences. These tRNAs are endogenously processed, allowing multiple gRNAs to be expressed from a single U6:3 promoter. This multiplexing strategy enhances cleavage efficiency at the target loci and minimizes formation of functional resistance alleles.

The first construct, designated as the *h*-drive (Figure 2A), is inserted into the endogenous *hairy* locus. To facilitate the identification of transgenic individuals, this cassette utilizes a 3xP3 promoter driving the expression of a DsRed fluorescent marker, enabling eye-specific screening. The core functional components include a *nanos* promoter-driven Cas9 nuclease and a U6:3 promoter-driven gRNA array. Crucially, these gRNAs are designed to act in *trans*, specifically targeting the wild-type *sim* gene. To maintain the viability of the carrier, the construct contains a recoded version of the *hairy* coding sequence followed by its natural 3’ UTR. This “antidote” sequence was engineered by synonymous codon replacement to remove all gRNA target sites and minimize nucleotide homology with the wild-type allele, thereby preventing self-cleavage and reducing the likelihood of genomic instability.

The second construct, designated as the *sim*-drive (Figure 2A), is inserted into the endogenous *sim* locus and carries a 3xP3-EGFP marker for distinct visual identification. This allele provides a recoded *sim* rescue sequence to protect its own function. Unlike a fully symmetric two-locus TARE design, however, the *sim*-drive uses an alternate gRNA strategy against *hairy*. Two of the gRNAs target distal *hairy* sequences and function as conventional TARE gRNAs, generating loss-of-function *hairy* alleles through end-joining or HDR. The second pair targets the region corresponding to the *h*-drive insertion site. Cleavage at this insertion-proximal region creates a potential opportunity for the wild-type *hairy* allele to use an *h*-drive-bearing homolog as a repair template, resulting in rare HDR-mediated copying of the *h*-drive (likely only when these gRNAs cut without the other pair cutting). Therefore, the *sim*-drive is designed both to disrupt wild-type *hairy* alleles and, in rare cases, to amplify *h*-drive alleles through HDR.

This asymmetric design allowed us to incorporate HDR-mediated drive conversion as a threshold-modulating process rather than as the primary drive mechanism. In the linked two-locus architecture, recombination and HDR are expected to act in opposite directions. Recombination breaks the linkage between different drive alleles, thereby increasing the introduction threshold. Rare HDR-mediated copying can amplify *h*-drive alleles, thereby lowering the threshold and partially counteracting recombination-mediated allele separation and the expected fitness cost at this site(Hou et al., 2024; Metzloff et al., 2022; Xu et al., 2026). Consistent with this logic, model scans showed that increasing drive conversion reduced the introduction threshold, whereas increasing recombination increased the threshold (Figure 2C). Thus, this system is designed as an asymmetric HDR-assisted linked two-locus TARE drive, in which linkage, sex-specific recombination, and HDR jointly tune the release threshold.

To experimentally parameterize the recombination component of this model, we measured the recombination rate between the *hairy* and *sim* loci using heterozygotes for two non-driving DsRed marker lines at these sites, one on each chromosome. Non-fluorescent progeny represented recombinant chromosomes lacking both DsRed markers. The total recombination rate was therefore calculated as twice the proportion of non-fluorescent progeny. This assay showed that recombination between *hairy* and *sim* was sex-specific, with no detectable recombination in males as expected and a female recombination rate of 30.6% (Data Set S3). These experimentally estimated values were used in the linked-drive model.

### Assessment of single-drive functionality

To assess the efficiency of the two separated drives, we first check the egg viability of drive crosses. We crossed drive heterozygotes and *w^1118^*wild-type flies for both the *hairy* and *sim* target loci. Eggs were counted, and adult survival was recorded to calculate egg-to-adult viability (Data Set S4). When female drive heterozygotes were crossed with *w^1118^* males, progeny viability was dramatically reduced (Figure 3A). For the *sim* drive, egg viability was 12.8% with a 95% CI of 5.7-26.4%, significantly lower than the *w^1118^* control (93.1%, 85.4-96.9%; *P* < 0.0001, z-test). For the *hairy* drive, viability was further reduced to 2.0% with a 95% CI of 0.7-5.3% (*P* < 0.0001, z-test). These results are consistent with the expected mechanism of the TARE drive with high but not 100% rates of cleavage from maternally deposited Cas9 and gRNA (Figure 3B). Because no progeny could inherit a drive allele at the cleaved locus in these female single-drive crosses, few offspring survived. Male drive heterozygotes likely had high germline cleavage rates due to the use of the *nanos* promoter (Carballar-Lejarazú et al., 2020; Du et al., 2024), which produces similar or even higher cleavage in females than males, and higher germline cleavage rates than embryo cleavage rates. However, because males do not deposit high levels of Cas9 into embryos, they produced progeny with viability comparable to wild-type controls, as expected from a target locus that is haplosufficient. Together, these results indicate that our target genes behaved as expected and that the toxin element of each drive functioned with high efficiency.

**Figure 3.**
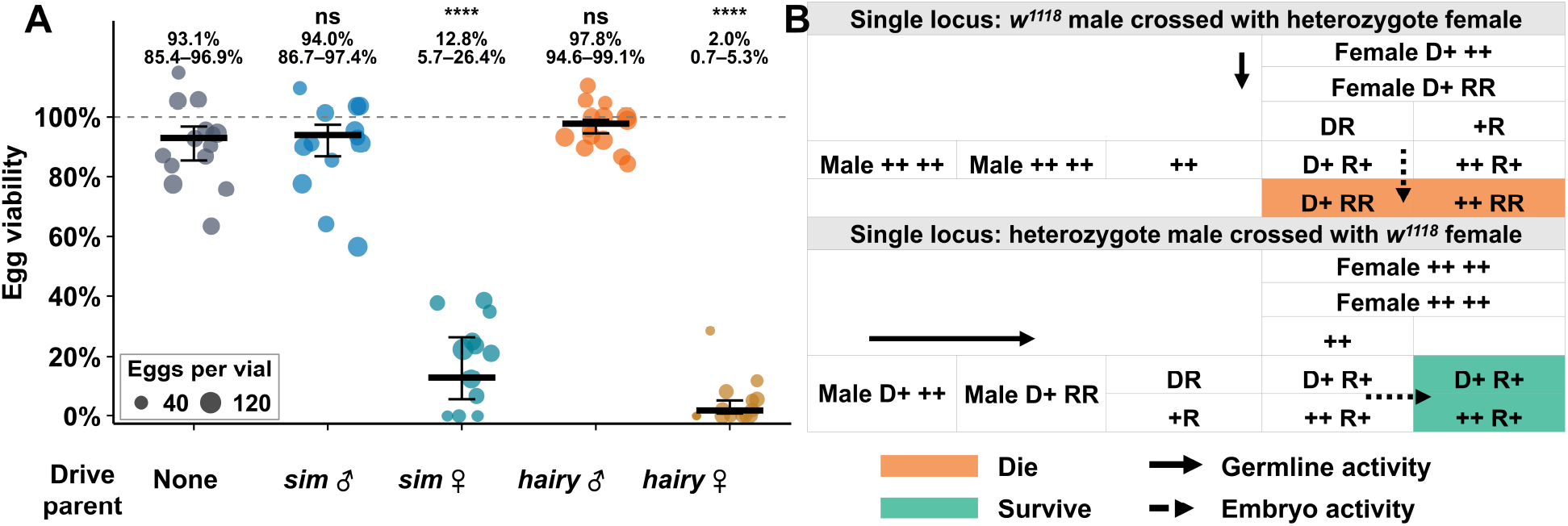
Single-locus drive performance. **(A)** Single-locus drive heterozygotes were crossed to *w^1118^* flies, and egg-to-adult viability was measured. Crosses are categorized by the drive-carrying parent, and “none” refers to control crosses where both parents were *w^1118^*. Each data point represents the result from a single vial. Horizontal lines indicate mean viability, and error bars indicate 95% confidence intervals. Values above each group indicate the model-estimated mean and 95% confidence interval. Significance symbols indicate comparisons to the *w^1118^* control (ns *P* ≥ 0.05, **** *P* < 0.0001). **(B)** Schematic illustration of expected inheritance patterns and viability outcomes for the male or female drive crosses. D denotes the drive allele, + denotes the wild-type allele, and R denotes the disrupted allele. Green shading indicates the genotypes of surviving progeny, while orange shading indicates progeny genotypes that are nonviable. Solid arrows represent Cas9 activity in the germline, and dashed arrows represent maternal Cas9 activity in the embryo.

### Evaluation of 2-locus TARE drive performance

To evaluate the combined activity of the 2-locus TARE drive system, we performed crosses between drive heterozygotes (carrying both *sim* and *hairy* drive alleles) and *w^1118^* flies, and we assessed both egg-to-adult viability and drive inheritance among progeny (Data Set S5). We found that the progeny of these double-heterozygote females exhibited highly biased inheritance for both loci (Figure 4A). The *sim* drive allele was inherited by 99.0% of the progeny, with a 95% CI of 97.5-99.6%, while the *hairy* drive allele was observed in 81.7% of offspring, with a 95% CI of 77.3-85.3%. Both values represented a significant deviation from the expected 50% Mendelian inheritance for crosses with one drive parent (*P* < 0.0001, z-test). The high drive inheritance rates observed were driven by substantial viability costs imposed on progeny that failed to inherit the drive alleles. In the cross between double-heterozygous females and wild-type males, egg-to-adult viability was drastically reduced to 21.1% with a 95% CI of 13.4-31.6% (Figure 4A, S1A), significantly lower than the 96.4% with a 95% CI of 92.0–98.5% observed in the *w^1118^* control (*P* < 0.0001, z-test). This result aligns with the expected mechanism where maternally deposited Cas9 disrupts the remaining paternal wild-type alleles at the essential loci (Figure 4B). The near-fixation of the *sim*-drive suggests highly efficient cleavage at that locus by the *hairy* drive, while the *hairy* locus showed slightly lower inheritance efficiency, suggesting a small fraction of wild-type alleles may have escaped cleavage or remained functional. In contrast, the progeny of males that were heterozygous for both drive alleles crossed to wild-type females did not show significantly altered inheritance for the *sim* drive (52.0%, 48.6-55.4%), while the *hairy* drive showed a modest but significant increase above Mendelian inheritance (56.4%, 53.0–59.8%), and progeny maintained high egg-to-adult viability (88.8%, 81.8-93.4%).

**Figure 4.**
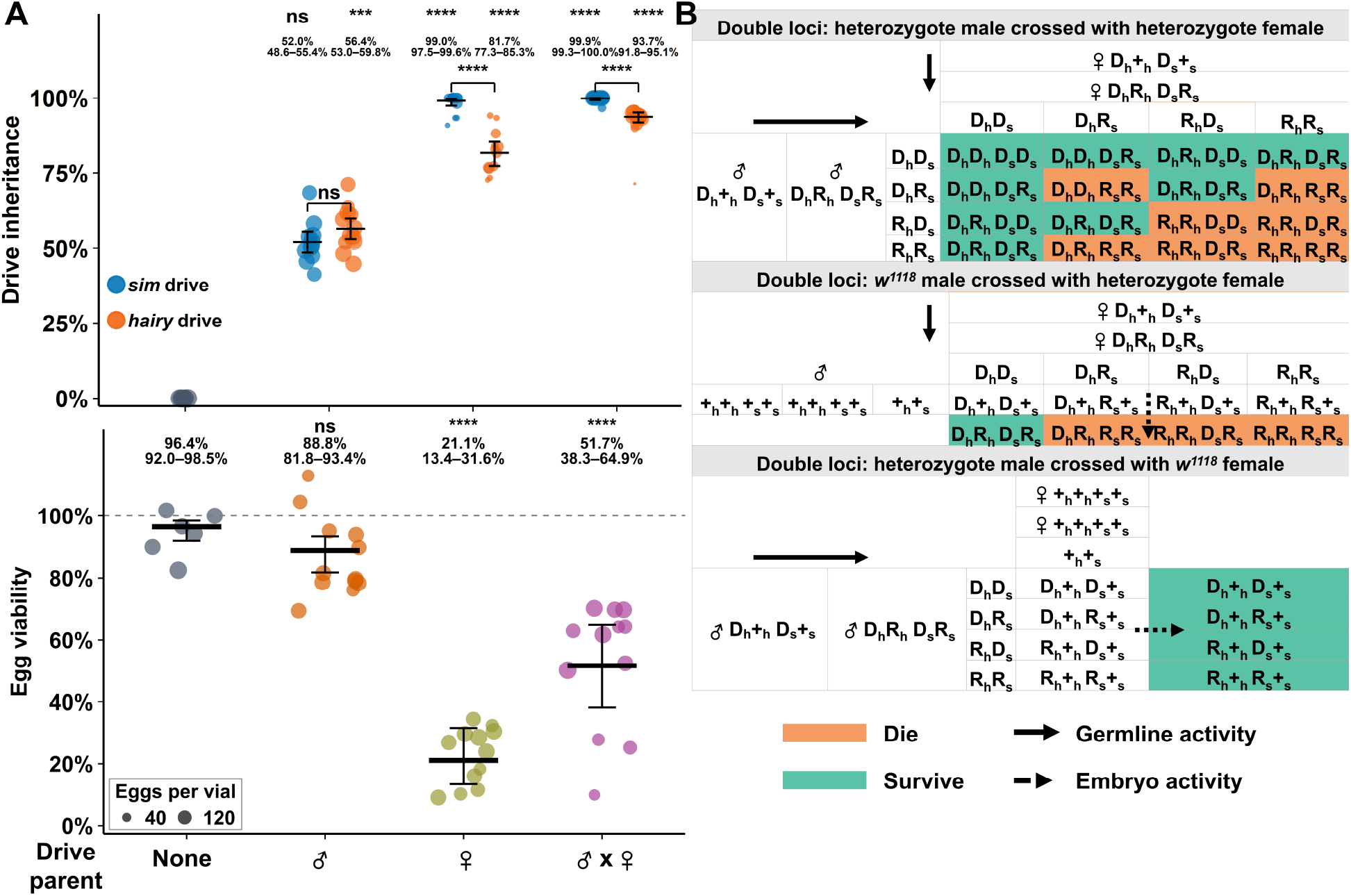
Performance of the 2-locus TARE drive system. **(A)** Crosses were performed using *w^1118^* flies and flies heterozygous for 2-locus TARE drive alleles at both the *sim* and *hairy* loci. Crosses are categorized by the drive-carrying parent(s), and “none” refers to control crosses where both parents were *w^1118^*. Each data point represents the result from a single vial. Horizontal lines indicate mean viability or drive inheritance, and error bars indicate 95% confidence intervals. Values above each group indicate the model-estimated mean and 95% confidence interval. Significance symbols indicate comparisons of drive inheritance against the expected Mendelian inheritance level, 50% for crosses with one drive parent and 75% for crosses with two drive parents, or of egg viability against the *w^1118^* control (ns *P* ≥ 0.05, *** *P* < 0.001, **** *P* < 0.0001, z-test). Note that egg viability higher than 100% represents undercounting of eggs in vials. **(B)** Schematic illustration of expected inheritance patterns and viability outcomes for each type of drive cross. Genotypes are indicated for both the *hairy* (h) and *sim* (s) loci: D denotes the drive allele, + denotes the wild-type allele, and R denotes the disrupted allele. Green shading indicates surviving genotypes, while orange shading indicates nonviable genotypes. Solid arrows represent Cas9 activity in the germline, and dashed arrows represent maternal Cas9 activity in the embryo.

Crosses between double-heterozygous females and double-heterozygous males led to an intermediate viability phenotype (51.7%, 38.3-64.9%). This matches the theoretical prediction that 9/16 (56.25%, assuming complete germline cleavage and no fitness costs) of progeny will survive by inheriting at least one rescue drive allele at each locus. Consequently, observed drive inheritance rates were extremely high at both loci (*sim*: 99.9%, 99.3-100.0%; *hairy*: 93.7%, 91.8-95.1%). Both values were significantly higher than the expected 75% inheritance for crosses with two drive parents (*P* < 0.0001, z-test). Theoretically, if germline cleavage efficiency is 100%, gametes will carry only drive or disrupted alleles, and maternal deposition is redundant, though it practice it can partially compensate for imperfect germline (Figure 4B). These results demonstrate that the double-locus TARE drive system functions as predicted in terms of both rescue element efficiency and toxin effect, exhibiting lethality for non-drive progeny and robust super-Mendelian inheritance for certain individual crosses, thereby showing potential for population modification.

To evaluate the trans-cleavage efficiency of the two spatially separated drive components in our system, we calculated the germline cleavage rates based on egg viability assays derived from crosses between single- or double-locus drive lines and wild-type individuals, as well as drive self-crosses. As summarized in Table 1, both the *h*-drive (targeting the distant wild-type *sim* locus) and the *sim*-drive (targeting the distant wild-type *hairy* locus) exhibited a germline cleavage rate of ∼1. This indicates highly efficient recognition and targeting of the distant alleles during gametogenesis. Furthermore, we observed substantial embryonic cleavage resulting from the maternal deposition of Cas9 and guide RNAs. Specifically, the maternal carryover of the *h*-drive machinery yielded a 97% embryonic cleavage rate at the *sim* locus (cleavage by the *hairy* drive), whereas the *sim*-drive yielded an 86% embryonic cleavage rate at the *hairy* locus.

**Table 1.** Germline and embryo cleavage rates from both drive elements.

| Locus | Target gene | Germline cleavage rate ($C_g$) | Embryo cleavage rate ($C_e$) |
| --- | --- | --- | --- |
| *h*-drive | *single-minded* | 1 | 0.97 |
| *sim*-drive | *hairy* | 0.97 | 0.86 |

### 2-locus TARE drive cage study

To assess the long-term population dynamics and invasion threshold of the 2-locus TARE system, several cage populations were established with varying initial frequencies of double drive double homozygotes. To initiate the cages, we released two groups of inseminated females: homozygous drive females (mated with drive homozygous males) and *w^1118^* females (mated with *w^1118^* males). These females were allowed to oviposit in food bottles for 24 hours before being removed. The release numbers were calibrated to achieve specific starting ratios, and their offspring were designated as “generation zero”. Based on the phenotypic screening of generation 0, the initial carrier frequencies (defined by the presence of both fluorescence markers) were 47%, 40%, 28%, 25%, and 9%. The cages were terminated when all individuals were carriers for both drive alleles (which also indicates that the presence of functional resistance is zero or very low) (Champer et al., 2020c). Alternatively, cages were stopped when the double drive carrier frequency dropped to 9%, which is far below even the ideal threshold and would soon lead to drive extinction. Throughout the study, the populations were maintained in discrete generations (Data Set S6).

In the two replicates initiated above the empirical threshold (yellow: 47%, orange: 40%), the 2-locus TARE system exhibited a steady increase in carrier frequency, reaching 100% by generation 16 and 20, respectively. This outcome demonstrates that when the release frequency surpasses a critical threshold, the drive’s inheritance bias effectively leads to successful population modification. Conversely, in the three replicates initiated at lower frequencies, the drive was progressively eliminated. In cages starting at intermediate frequencies (brown: 28%, blue: 25%) and low frequency (green: 9%), the carrier frequency steadily declined and failed to establish (Figure 5A-B, S2A-E). While the theoretical introduction frequency threshold for an ideal 2-locus TARE system is approximately 18% in the absence of fitness costs, the failure of the 25% and 28% releases indicates a substantial fitness cost associated with the drive alleles in this specific genomic context. This fitness burden imposes a selective disadvantage that requires a higher initial frequency to overcome, effectively raising the empirical threshold between approximately 28% and 40%.

**Figure 5.**
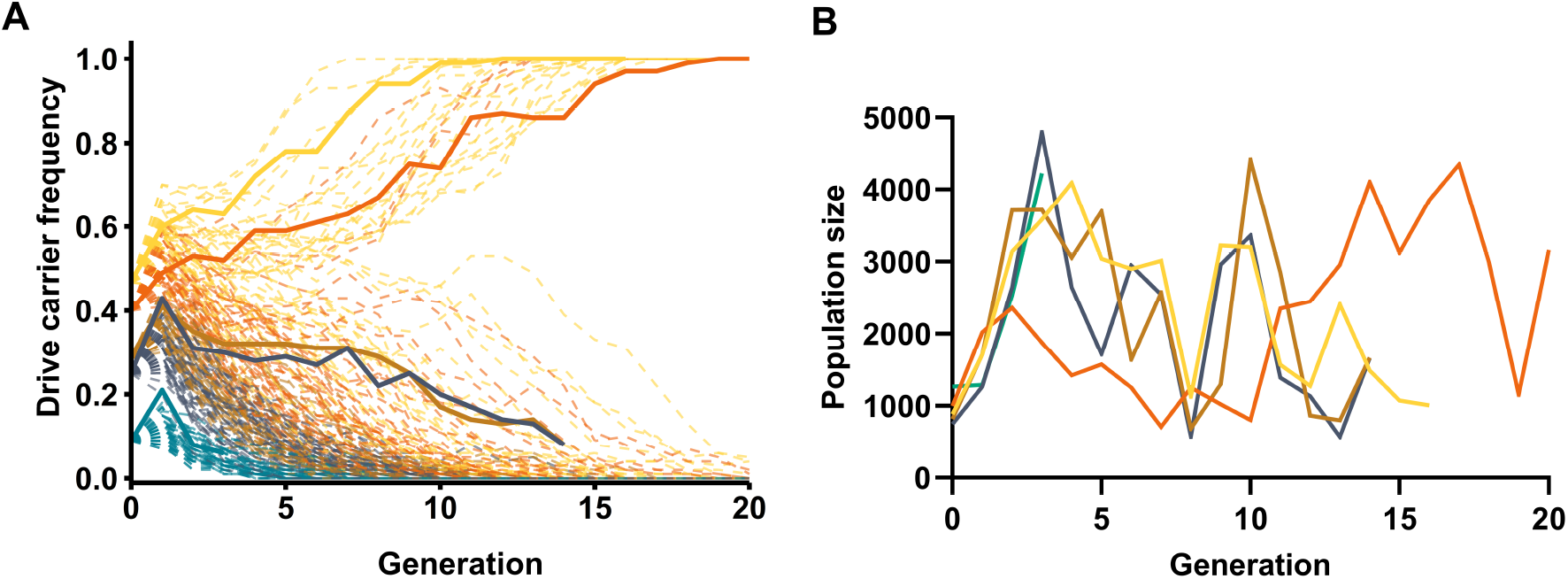
Performance of the 2-locus TARE drive in population cages. Cage trials were established by mixing double homozygous drive flies with age-matched *w^1118^* flies. Populations were maintained in enclosures with discrete generations (12-14 days per cycle), where adult flies were phenotyped and removed after a 24-48 hour egg-laying period to seed the subsequent generation. **(A)** Drive carrier frequency was determined by the proportion of individuals expressing both EGFP and DsRed fluorescence markers. Model populations were initiated with the same starting frequencies of 2-locus TARE drive homozygotes as in the cages: 47%, 40%, 28%, 25%, and 9% (each a different color). Dashed lines show simulated cage populations by R using parameters derived from experimental data. All simulations were performed assuming an effective population size of 100 with 20 replicates per starting frequency. **(B)** Adult population size was monitored across for each cage.

### Maximum likelihood analysis of drive fitness

To evaluate the fitness of the drive alleles in our cage population studies, we applied a maximum likelihood approach utilizing the germline and embryonic cleavage rates derived from our experimental egg-to-adult viability assays (Table 1). We inferred the fitness of each drive allele for drive homozygotes assuming a codominant model with multiplicative fitness per allele, where a value of 1 is equivalent to wild-type fitness. To determine model quality and select the most parsimonious fit for the observed cage data, we used Log-likelihood (logL) to indicate relative probability and the corrected Akaike Information Criterion (AICc) to penalize overfitting. Table 2 shows that the inferred fitness for the *hairy*-drive was significantly lower than wild-type (0.63), consistent with previous studies involving *hairy* drive modifications in other species (Hou et al., 2024; Metzloff et al., 2022; Xu et al., 2026). In contrast, the *sim*-drive fitness estimate was comparable to wild-type, indicating that our recoded *sim* sequence successfully rescued the essential gene function and had no unexpected off-target effects. These findings highlight *sim* as a potentially superior target for essential, haplosufficient gene drives due to its high cleavage susceptibility (though our increased cleavage rates could potentially be due to specific gRNA sites and expression from the *hairy* site) and robust functional rescue. Indeed, the frequency of individuals carrying the sim-drive increased a little more rapidly (Figure S2A).

**Table 2.** Maximum likelihood fitness parameter estimates from cage populations.

| Fitness costs | logL value | AICc | Ne | *hairy*-drive viability fitness | *sim*-drive viability fitness |
| --- | --- | --- | --- | --- | --- |
| None | -10.49 | 23 | 2.51 [2.51, 3.27] | 1 | 1 |
| *hairy* drive | -6.35 | 16.9 | 2.82 [2.82, 3.59] | 0.63 [0.45, 0.86] | 1 |
| *sim* drive | -8.47 | 21.1 | 2.65 [2.65, 3.42] | 1 | 0.72 [0.52, 0.99] |
| *hairy* drive;<br>*sim* drive | -6.35 | 19.1 | 2.82 [2.82, 3.59] | 0.63 [0.45, 0.86] | 1.00 [0.77, 1.00] |
| *hairy* drive =<br>*sim* drive | -7.21 | 18.6 | 2.75 [2.75, 3.51] | 0.81 [0.68, 0.95] | 0.81 [0.68, 0.95] |

Values in brackets denote the 95% confidence intervals. logL - log(likelihood), Ne - effective population size, AICc - Akaike Information Criterion (corrected).

All models additionally incorporated an effective population size (ne). Notably, our estimate for the effective population size was low, potentially indicating a poor model fit, despite the high confidence of the fitness cost in *hairy*. This discrepancy likely stems from significant fluctuations in population dynamics. While the populations were initially large, the cages suffered a fungal or bacterial infection mid-experiment, which caused substantial fluctuations in the life cycle and mortality rates of the flies that may have affected the gene drive, perhaps increasing or reducing its fitness under these conditions (especially in the three intermediate release level cages).

We validated the model by replicating the release conditions used in the physical cage experiments with a low effective population size (Figure 5A). The simulations broadly recapitulated the bistable dynamics observed in the cage experiments: releases initiated at higher frequencies (47% and 40%) could potentially result in the spread of the carrier allele toward fixation, whereas releases initiated at lower frequencies (28%, 25%, and 9%) were followed by its loss. This pattern is consistent with the relatively small effective population sizes inferred from the cage experiments by maximum likelihood estimation and suggests that drift or other factors contributed substantially to the observed fluctuations while preserving the overall bistable behavior.

To determine the introduction threshold of the 2-locus TARE drive system under the current parameter settings, we performed a systematic scan of introduction frequencies, which indicated that the introduction threshold was close to 50% (Figure S3A, Data Set S7).

## Discussion

Our study successfully demonstrates the design and empirical validation of an asymmetric linked 2-locus TARE gene drive in *Drosophila melanogaster*. Experimental results, ranging from individual crosses to multigenerational cage populations, align closely with the system’s theoretical framework. Its primary strategic advantage over standard TARE drive is the existence of a predictable introduction threshold. This property creates a bistable dynamic: when the frequency of drive-carrying individuals is below this critical threshold, natural selection favors wild-type alleles; conversely, when the release frequency surpasses this threshold, the drive spreads. This inherent threshold provides a crucial layer of ecological safety, creating a system that is potentially spatially confinable. It mitigates the risk of uncontrolled spread from accidental or small-scale releases because the drive cannot establish from a low frequency. Our cage studies empirically confirmed this threshold, locating it approximately between 28% and 40% (perhaps as high as 50% based on model analysis) for our specific constructs, a value elevated from the ideal theoretical prediction of 18% due to observable fitness costs, despite the genetic architecture of the linked two-locus system. This confirmation of a robust threshold demonstrates the potential of the 2-locus TARE system for localized and reversible population modification applications.

As engineered, 2-locus TARE functions as a modification drive designed to replace a wild-type population with one carrying a desired genetic trait. For a 2-locus TARE system, there are two primary strategies for cargo gene integration: the cargo can be inserted into both drive units or restricted to a single locus. While dual-locus insertion theoretically expands the reach and redundancy of the cargo gene, single-locus insertion offers distinct advantages by minimizing the additional fitness costs often associated with large genetic payloads, as well as simplifying engineering. In our experimental evaluation, potentially integrating the cargo into the *sim*-drive locus emerged as a viable option. Compared with using both loci for cargo integration (Figure S2B), utilizing this higher-performance locus exclusively for cargo could therefore represent a feasible design choice, because it may preserve modification potential while reducing construct size and cargo-associated fitness costs. However, future modeling will be required to determine whether single-locus cargo insertion indeed provides superior overall performance, particularly because cargo retention and spread may depend on linkage, recombination between the two drive loci, and the relative fitness costs of the two drive components.

As with the standard TARE drive, 2-locus TARE cannot be effectively converted into a suppression drive on its own. This is because both 2-locus TARE and standard TARE drives require a prolonged period to accumulate drive homozygotes. Consequently, they generate insufficient genetic load to eliminate a population. Nevertheless, a tethered drive strategy can overcome this limitation (Dhole et al., 2019; Feng and Champer, 2024; Metzloff et al., 2022). By using a fully functional TARE or 2-locus TARE drive as a “tether” to confine the release area of a split homing suppression drive, confined population suppression can be achieved. This approach allows the system to utilize the suppression power of homing drives while fully using the inherent confinement characteristics of the toxin-antidote architecture. However, to avoid failure from simultaneous or closely timed releases, it is important to ensure that the 2-locus TARE drive spreads sufficiently in space before the suppression drive arrives (Feng and Champer, 2024).

While our cage study provided essential empirical validation of the introduction threshold in a panmictic setting, a drive’s true confinement potential must be assessed in realistic populations, as spatial dynamics can profoundly alter drive behavior and lead to divergent outcomes between different modeling frameworks. Modeling based on linked-deme frameworks, where discrete populations are connected by a defined migration rate, predicts effective confinement for certain real-world populations. However, the invasive potential of a drive, is highly dependent on the population’s spatial structure. In many scenarios, spatially continuous populations may be more representative. In these, a similar 2-locus underdominance system showed high invasive potential (Champer et al., 2020d). The discrepancy arises from the drive’s ability to propagate as a wave in a cohesive front, maintaining the high local allele frequencies needed to fuel its advance. The 2-locus TARE system’s low introduction threshold (<50%) would also enable such a self-propagating wave to form and advance into wild-type territory from a sufficiently large and dense release area. Some obstacles may be able to stop the wave of advance, but these will need to be considered and modeled for any specific deployment where confinement is desired.

Our empirical data indicate that drive-associated fitness costs elevated the observed introduction threshold well above the ideal level. This highlights the necessity of minimizing design-imposed fitness burdens. In our 2-locus TARE system, the observed fitness costs likely stem from both the fundamental mechanism of the toxin-antidote principle and the structural design of the 2-locus framework. First, regarding the toxin-antidote mechanism, utilizing a same-site rescue strategy avoids the position effects that can be associated with distant-site rescue (Chen et al., 2023). However, the modification may still disrupt native regulatory architectures such as cryptic enhancers residing within removed introns. Furthermore, the recoded rescue element might suffer from suboptimal translation efficiency if codon usage was altered too drastically to without rigorous species-specific optimization Second, concerning the 2-locus framework, the integration of two independent Cas9 expression cassettes imposes a continuous metabolic load on the host and exacerbates global off-target toxicity, which appears to have measurable fitness effects in at least some situations (Langmüller et al., 2022). Additionally, the presence of large stretches of identical sequences across the two loci drastically increases the risk of ectopic homologous recombination. These identical sequences include the duplicated promoters, Cas9 coding regions, and terminators, which together could lead to potential genomic instability. It also remains possible that the target genes were not entirely haplosufficient.

To mitigate these challenges and improve overall drive performance, future designs should incorporate several structural refinements. Addressing the translational and regulatory defects within the recoded rescue element requires an advanced context-aware codon optimization strategy. Recent genome-wide translatomic analyses in *Drosophila melanogaster* demonstrate that optimal codons not only facilitate faster ribosomal elongation but also significantly reduce amino acid misincorporation rates (Wu et al., 2024). This underlying biological dynamic necessitates a refined recoding scheme capable of precisely balancing the reduction of sequence homology with the preservation of translational efficiency. Similarly, the structural complications arising from the 2-locus framework can be resolved by eliminating genetic redundancy. Employing an alternate Cas9 sequence at the second locus would help remove sequence homology within the nuclease open reading frames, and different Cas9 promoters with similar efficiency could further reduce identity (Du et al., 2024; Wu et al., 2025).

However, the efficiency and suitability of promoters is highly context-dependent and varies between species. For instance, the first homing drive in *Anopheles stephensi* using a *vasa*-Cas9 promoter displayed an intermediate level of embryo resistance (Gantz et al., 2015). In *Anopheles gambiae*, promoters such as *nanos* and *zpg* markedly reduced negative somatic effects and achieved much lower rates of embryo resistance while retaining high drive efficiency (Hammond et al., 2021). In *Aedes aegypti*, even the most effective homing drives utilize promoters that are not completely restricted to the germline (Anderson et al., 2023; Anderson et al., 2024). Ultimately, this highlights that each species possesses a unique toolkit of effective promoters, dictating whether a strategy prioritizing a low invasion threshold or one aiming for rapid population modification is not only optimal, but biologically feasible.

To conclude, we successfully validated the 2-locus TARE system in *Drosophila melanogaster* using a combination of essential, haplosufficient genes. This architecture allows for efficient self-propagation while maintaining a safety-conferring introduction threshold. By effectively coordinating self-rescue with distant-site targeting, the system offers a robust, versatile template for localized and reversible population modification in non-model organisms. Beyond the drive mechanism itself, our findings confirm that *single-minded* (*sim*) serves as a good target locus, with minimal fitness costs affecting the drive at this site. We demonstrated that *sim* is both essential and haplosufficient, characterized by high cleavage susceptibility and robust functional rescue efficiency. Due to its high level of sequence conservation, *sim* is also a promising candidate for expansion to other species.

## Acknowledgements

The cluster-based data collection was assisted by High-Performance Computing Platform of the Center for Life Sciences at Peking University. This study was supported by the Center for Life Sciences and the National Natural Science Foundation of China (grants 32270672 and W2432018).

## Supplementary Information

**Figure S1.**
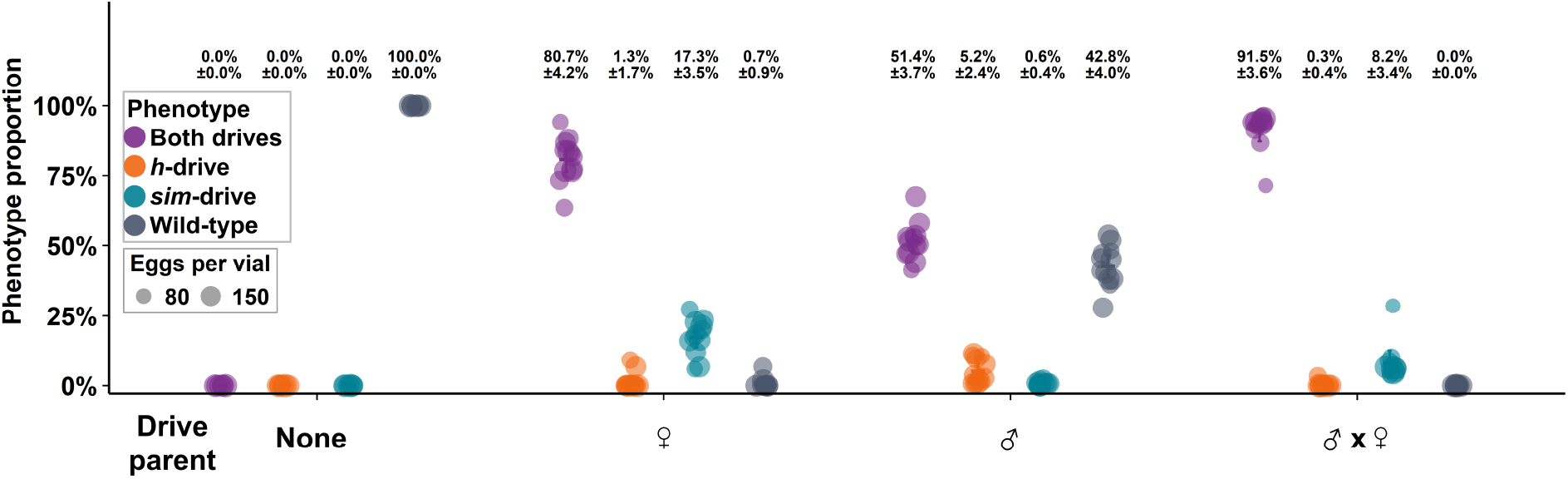
Progeny phenotype distributions from individual crosses. Crosses were performed using *w^1118^* flies and flies heterozygous for 2-locus TARE drive alleles at both the *sim* and *hairy* loci. Crosses are categorized by the drive-carrying parent(s), and “none” refers to control crosses where both parents were *w^1118^*. Each data point represents the observed proportion of a specific offspring phenotype from a single vial. Horizontal lines indicate mean phenotypic frequencies, and error bars indicate 95% confidence intervals.

**Figure S2.**
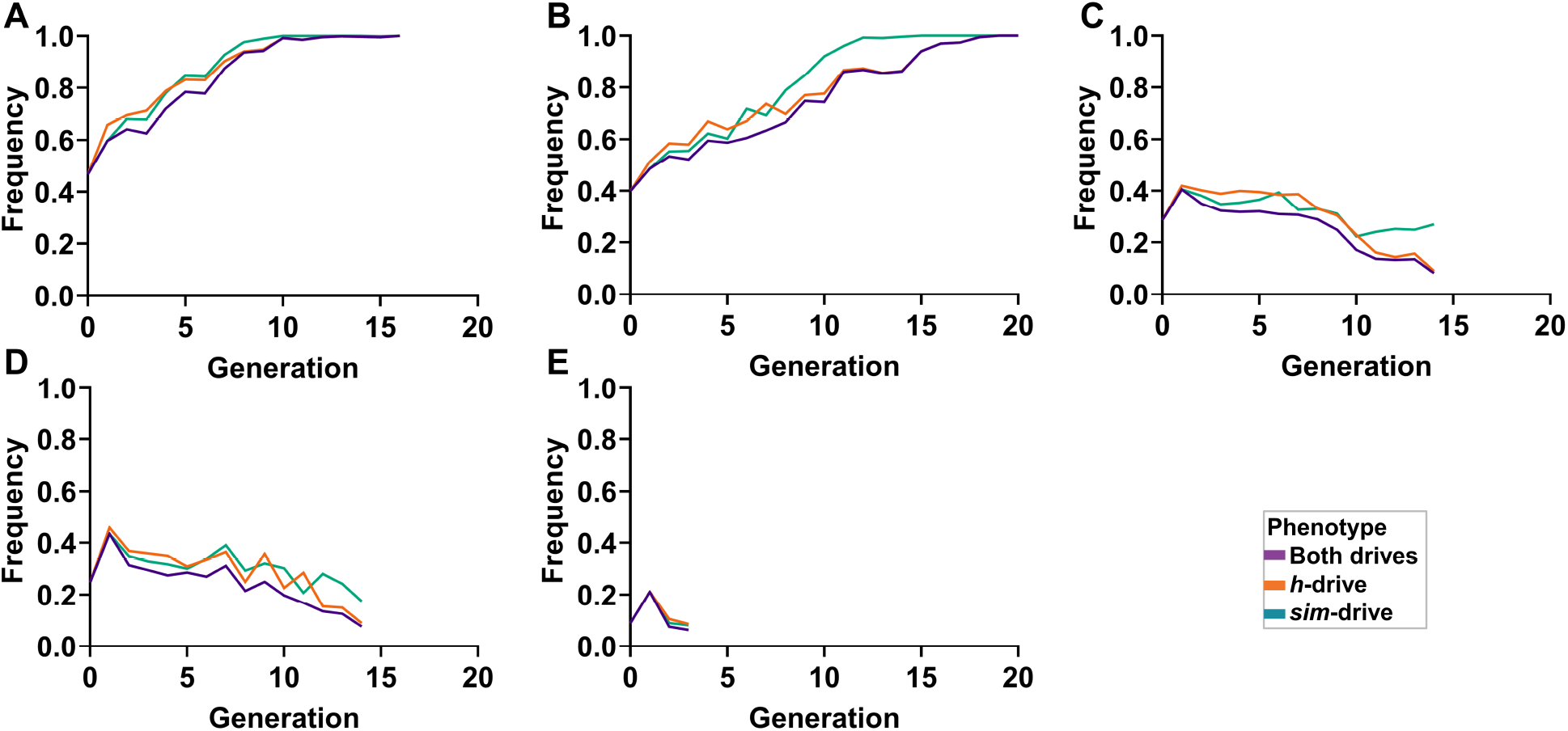
Additional details of the 2-locus TARE drive in population cages. (A-E) Frequencies of both-drive, h-drive, and sim-drive phenotypes were tracked across generations in five independent cage populations with different initial both-drive frequencies. Purple, orange, and green lines represent both drives, *h*-drive, and *sim*-drive, respectively. Cages initiated at high both-drive frequencies increased toward fixation, whereas cages initiated below the threshold declined, indicating bistable population dynamics. The divergence between successful and unsuccessful cages supports the presence of an introduction threshold for the linked two-locus TARE drive.

**Figure S3.**
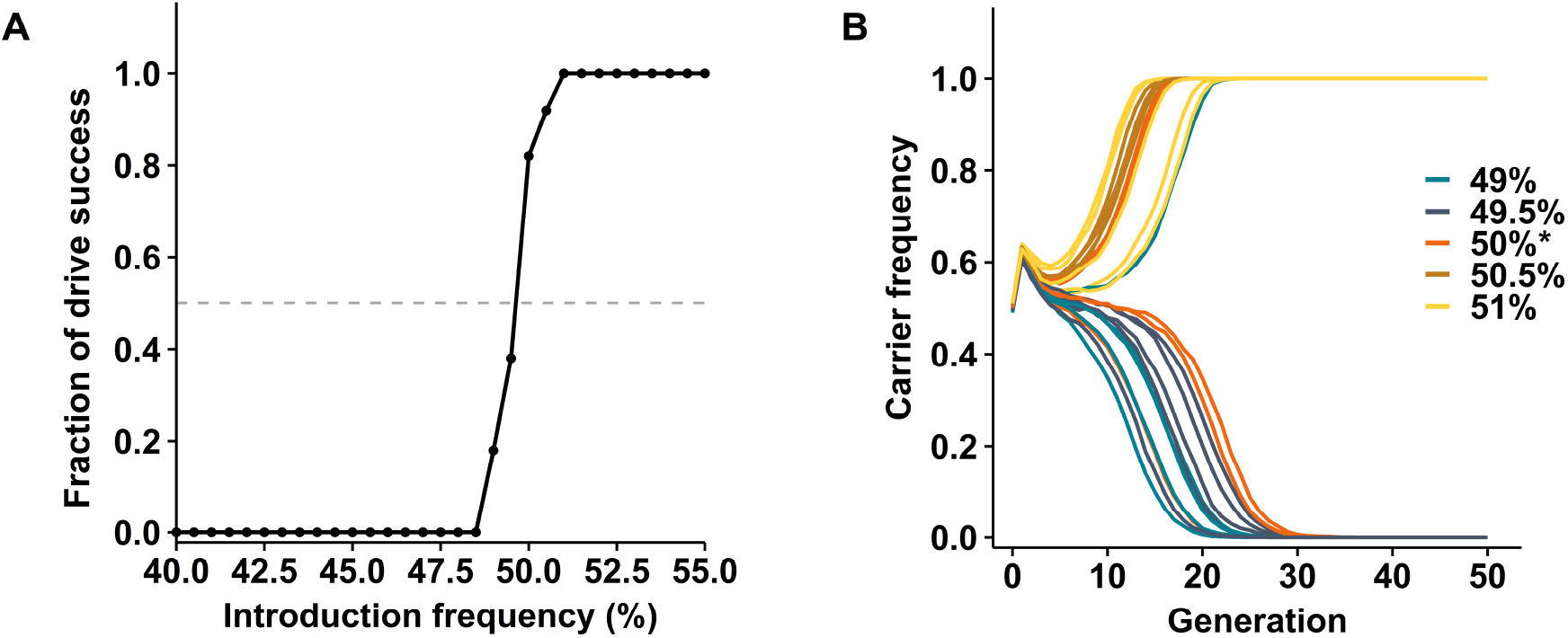
Estimation of the introduction threshold for the 2-locus TARE gene drive system. **(A)** The proportion of simulation replicates in which the drive successfully established is plotted against the introduction frequency of drive homozygotes. Drive establishment was defined as a strict, consecutive increase in total carrier frequency for at least 10 generations within the first 50 generations post-release. At each frequency, 20 simulation replicates were performed. The dashed grey horizontal line indicates the 50% proportion criterion used to define the introduction threshold. **(B)** Initial release frequencies were varied in 1% increments (from 49% to 51%), using 5 replicates per frequency. * represents the predicted introduction threshold. All simulations were performed with an effective population size of 10,000 by R. Carrier frequency indicates individuals with at least one drive allele (of either type).

